# Adaptation-induced sharpening of orientation tuning curves in the mouse visual cortex

**DOI:** 10.1101/2023.12.24.573226

**Authors:** Afef Ouelhazi, Vishal Bharmauria, Stéphane Molotchnikoff

## Abstract

Orientation selectivity is an emergent property of visual neurons across species with columnar and non-columnar organization of the visual cortex. To compute the orientation selectivity of a neuron, a tuning function is fit on the raw responses of a neuron and then a measure, termed orientation selectivity index (OSI), is derived from this fitted curve to determine the tuning of the neuron. Previously, it has been shown that adaptation (a protocol where a neuron under observation is presented a non-optimal stimulus for a specific time) has varying effects on the tuning properties of neurons, such as, orientation, spatial frequency, motion etc. The emergence of OSI is more established in columnar cortices than the non-columnar ones. However, how adaptation impinges upon the OSI of the latter has not been systematically investigated. Here, in the mouse primary visual cortex (V1), we show that a 12-min adaptation protocol sharpens the OSI (tuning) of the visual neurons, underlying a specific dendritic neural mechanism, potentially facilitating the learning of novel features.

## 1. INTRODUCTION

To encode visual inputs, cortical neurons, embedded in a microcircuit, must generate selective responses for distinct stimulus features. In the primary visual cortex (V1), the selective responses of neurons to the orientation of edges have been extensively studied across species (Campbell et al., 1968; Carandini et al., 1998; Hubel and Wiesel, 1965, 1959; Kaschube et al., 2010; Monier et al., 2003; Tan et al., 2011; Wiesel and Hubel, 1963; Wilson et al., 2016). Many studies have shown that sharp selectivity is observed even in species where V1 lacks an orientation map, such as rodents (Bonin et al., 2011; Cossell et al., 2015; Jeyabalaratnam et al., 2013; Niell and Stryker, 2010; Ohki et al., 2005; Pattadkal et al., 2018; Weiler et al., 2022). To determine the orientation selectivity of a neuron, generally, an orientation selectvity index (OSI) is computed from the tuning function of the neuron (Bachatene et al., 2013; Bharmauria et al., 2016b; Mazurek et al., 2014; Niell and Stryker, 2008; Womelsdorf et al., 2012).

Adaptation, an experimental protocol where a non-preferred stimulus (adapter) is imposed on the neuron(s) under observation, can alter the orientation tuning of neuron along with their OSI (Bharmauria et al., 2022; Kohn, 2007; Tring et al., 2023). For example, adaptation can shift the peak of the orientation tuning curve either toward (attractive shift) the adapter or away (repulsive shift) from it (Bharmauria et al., 2022; Dragoi et al., 2000; Ghisovan et al., 2009; Kohn, 2007). Continuously presenting the optimal stimulus to the neuron leads to suppression of the response to that stimulus, thereby, broadening of the tuning curves (Giaschi et al., 1993; Harris et al., 2000; Kohn, 2007). Notably, the duration of adaptation, intensity of the stimulus, and other contextual factors may have distinct effects on neuronal properties (Bao et al., 2013; Bharmauria et al., 2022; Kohn, 2007; Roach and McGraw, 2009).

OSI is highly heterogeneous across species and neurons: V1 neurons of lower vertebrates are generally weakly tuned than V1 neurons of higher vertebrates (Davey et al., 2022; Nguyen and Freeman, 2019; Tan et al., 2011) and fast spiking neurons (putative inhibitor interneurons) mostly exhibit broader tuning than the regular spiking neurons (putative pyramidal cells) (Bharmauria et al., 2015; Chanauria et al., 2016; Insel and Barnes, 2015; Schneider et al., 2023). As V1 neurons in higher vertebrates are systematically arranged into orientation columns, the origin of OSI is more established in higher vertebrates than in species with salt-and-pepper organization (Alitto and Dan, 2010; Davey et al., 2022; Pattadkal et al., 2018; Tan et al., 2011; Tang et al., 2023).

What are the effects of the visual adaptation on mouse visual neurons and how does new selectivity emerge post-adaptation? Here, using a 12-min uninterrupted adaptation protocol (Afef et al., 2022; Bachatene et al., 2015b; Jeyabalaratnam et al., 2013) on mouse visual neurons, we examined the above questions. We report that adaptation further sharpens the selectivity of mouse visual neurons, suggesting emergence of novel circuits through changed excitation/inhibition balance. This sharpening of the tuning curve may facilitate learning of novel features.

## 2. MATERIALS AND METHODS

The animal studies were approved by the Institutional Animal Care and Use Committee of the University of Montreal following the guidelines of the Canadian Council on Animal Care. Experiments (animal surgery procedures and electrophysiological recordings) were performed using nineteen CD-1 strain adult mice (weight: 28-31□g, 9-11□weeks). Animals were supplied by the Division of Animal Resources of the University of Montreal. Anesthetized mice were prepared for electrophysiological recordings in the V1 as described below.

### 2.1 Animal preparation, visual stimulation, and electrophysiological recording

Mice were deeply anaesthetized with 10% urethane (1.5□g/kg), injected intraperitoneally. A local anesthetic [Lidocaine hydrochloride 2% (Xylocaine, AstraZeneca, Mississauga, ON, Canada)] was administered subcutaneously at the surgical and pressure sites at the start of the surgery. An incision was made into the skin over the brain and a dural flap was made (2.5 × 2.5 mm) over the visual cortex. Mice were placed in a stereotactic frame allowing monocular visual stimulation of the entire contralateral visual field. The unstimulated eye was closed. Tungsten microelectrodes (2–10 MΩ at 1 kHz; Frederick Haer & Co, Bowdoinham, ME, USA) were inserted at a depth less than 1 mm to record the signal. The latter was amplified, band-pass filtered (300 Hz–3 kHz), digitized and recorded with a data acquisition software (Spike 2, Cambridge Electronic Design, CED Limited, Cambridge, England).

### 2.2. Monocular stimulation

Drifting sine wave gratings generated with a VRG Volante 34020 graphic board (Vision Research Graphics, New Hampshire, USA) were presented, with an eye-to-screen distance of 30 cm, on a 21-inch monitor (60 Hz refresh rate, Mitsubishi FHS6115SLK Color Display Monitor, Tokyo, Japan) with 1024×512 pixels resolution. The center of the monitor was positioned at about 30.7° azimuth, 0° elevation. Spatial frequency, temporal frequency, and velocity were set to evoke optimal firing in mouse, and fixed at 0.07 cycles/°, 2 Hz, and 4°/s, respectively. Eight orientations covering a span of 157.5°, equally spaced and shown in a random order were presented in the control condition (**Fig .1A**). Each orientation was applied in blocks of 25 trials of 4.096 s each, separated by an interval of 1.0 - 3.0 s during which a dark screen was presented. An adaptation session followed the control phase where an adapting stimulus (90°) was presented continuously for 12 minutes. Test orientations were presented immediately following the adaptation period. Control and post-adaptation recordings were carried out, but no recordings were performed during adaptation session (**Fig. 1B**). Action potentials were sorted from the multi-unit signal using the spike sorting method and further confirmed by cluster analysis (Afef et al., 2022; Bharmauria et al., 2016a; Chanauria et al., 2019). The stability of each cell’s response across conditions was verified qualitatively by visual control of the cluster disposition and of the waveform shape.

**Fig 1.**
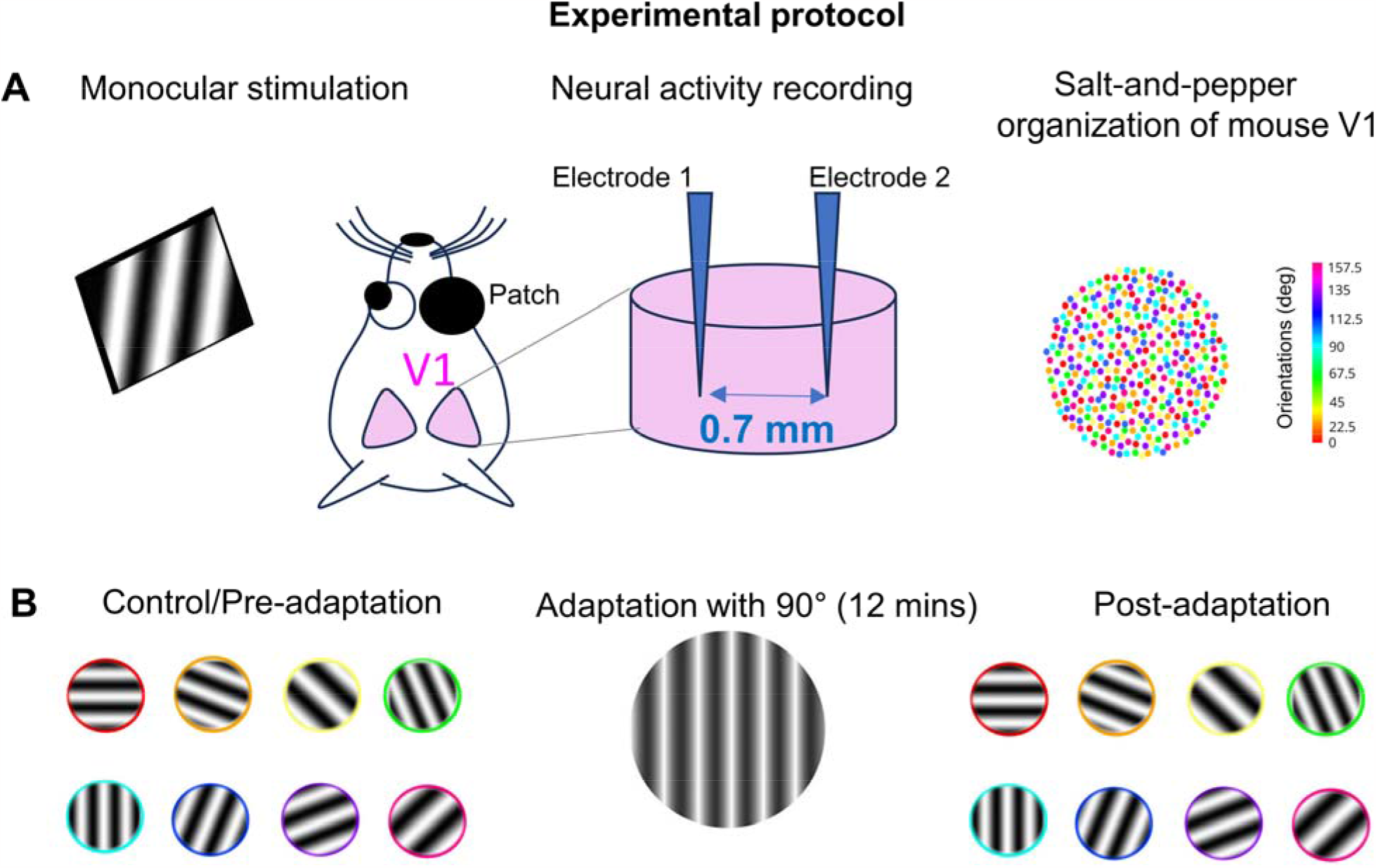
Experimental design. **(A)** Cartoon displaying visual stimulation (left) on the mouse striate cortex. Two multisite electrodes (Electrode 1 and Electrode 2, interelectrode distance = 0. 7 mm) were inserted into V1 (middle) to record cells with different preferred orientations (PO), organized in a salt-and-pepper fashion (right). **(B)** Experimental time course. Recordings were performed in the control condition to the orientation test set (left), and after 12 minutes (right) of uninterrupted presentation (middle) of an adaptor at 90°. All the orientations were randomly presented on the screen in 25 trials with arbitrary intervals (1-3 s).

Then, orientation tuning curves were constructed from the raw data and fitted with the von Mises function (Swindale, 1998) allowing a precise determination of the neuron’s preferred orientation (PO). The von Mises function is defined as:

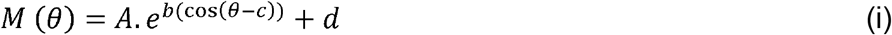

where A is the value of the function at the PO, c, and b is a width parameter. An additional parameter, d, represents the spontaneous firing rate of the cell (Kohn and Movshon, 2004; Swindale, 1998).

It was also necessary to determine the degree of tuning of each cell in our sample. An orientation selectivity index (OSI) was calculated to ensure the tuning of neurons. It was measured using the fitted tuning curves by dividing the firing rate at the baseline (orthogonal orientations) by the firing rate at the preferred orientation and subtracting the result from one. The closer the OSI is to one, the stronger the orientation selectivity (Bharmauria et al., 2016b; Liao et al., 2004; Ramoa et al., 2001). In this study, an OSI threshold of 0.4 was fixed for tuned cells (T) (Atallah et al., 2012; Bachatene et al., 2016). Therefore, cells with an OSI ≥ 0.4 were considered tuned and with and OSI < 0.4 were deemed untuned.

## 3. RESULTS

We elucidated the effect of adaptation on the OSI of mouse visual neurons pre- and post-adaptation. Out of 113 investigated cells in this study, 47 (41.23%) cells were selectively tuned (T) and 67 (58.77%) cells were untuned (U) in the control condition; consistent with the proportion in previous studies (Atallah et al., 2012; Bachatene et al., 2016; Jeyabalaratnam et al., 2013). In the following sections, we determine how the tuning properties of these two classes of cells change post-adaptation. This analysis is based on an OSI cut-off threshold of 0.4: neurons with an OSI of < 0.4 were considered untuned and > 0.4 were categorized as tuned neurons.

### 3.1. Effect of adaptation on the OSI of cells

We began by investigating the tuned population of cells in the control condition, i.e., how their OSI changed post-adaptation. 30 / 47 (63.2 %) retained their tuning, whereas 17 / 47 (36.17 %) became untuned. The same analysis for the untuned population revealed that most cells (52 / 67, 77.61 %) remained untuned but a proportion (15 / 67, 22.39 %) of cells exhibited tuning. Notably, following adaptation (**Fig. 2A**), similar proportions of tuned (39.47 %) and untuned cells (60.52 %) units were observed compared with the control condition (tuned = 41.23%; untuned = 58.77 %) (McNemar test, 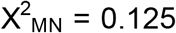, p = 0.75). Further, the OSIs of pre- (0.37 ± 0.21, mean ± SEM) and post-adaptation (0.36 ± 0.25) populations were not significantly different (Wilcoxon paired test, p = 0.96) from each other.

**Fig 2.**
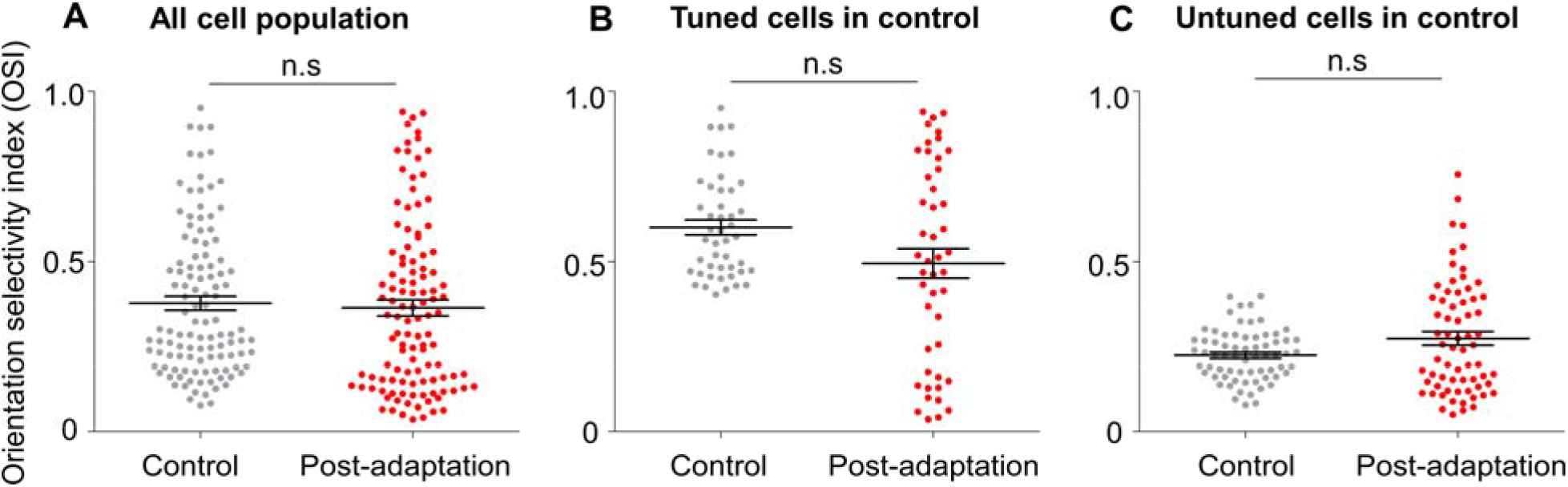
Population statistics. **(A)** No significant difference of OSI between the pre- and post-adaptation cell populations (tuned + untuned). **(B)** No significant difference in OSI of tuned cells between the pre- and post-adaptation. **(C)** No significant difference in OSI of untuned cells between pre- and post-adaptation.

When we performed this analysis for the tuned (**Fig.2B**, OSI pre-adaptation = 0.60±0.15, post-adaptation OSI = 0.49 ± 0.29; Wilcoxon paired test, p = 0.08) and untuned cells pre- and post-adaptation (**Fig. 2C**, control OSI = 0.22 ± 0.07, post-adaptation OSI = 0.27 ± 0.16; Wilcoxon paired test, p = 0.07), again, no significant difference was observed: Collectively, this suggests an occurrence of a homeostatic mechanism between pre- and post-adaptation neural circuits (Bachatene et al., 2015b; Benucci et al., 2013; Turrigiano and Nelson, 2000).

We then proceeded with in-depth analysis of these subpopulations on a pair-wise basis, i.e., tracking the OSI of a cell pre- and post-adaptation. The effect of adaptation on OSI values was different depending on the sub-populations (untuned-untuned, tuned-tuned, tuned-untuned and untuned-tuned). While the adaptation failed to change the OSI of untuned-untuned sub-population (**Fig. 3A**, Paired t-test, p = 0. 31), the OSI increased significantly in the tuned-tuned sub-population (**Fig. 3B**, Paired t-test, p = 0.009, t = 2.772, df =34). For the tuned-untuned sub-population, adaptation significantly decreased the OSI by 26.52% (**Fig. 3C**, Paired t-test, p < 0.0001, t = 7.713, df =15). By contrast, following adaptation, the untuned-tuned sub-population showed a significant increase of their OSI by 48.7% (**Fig. 3D**, Paired t-test, p < 0.0001, t = 9.193, df =16).

**Fig 3:**
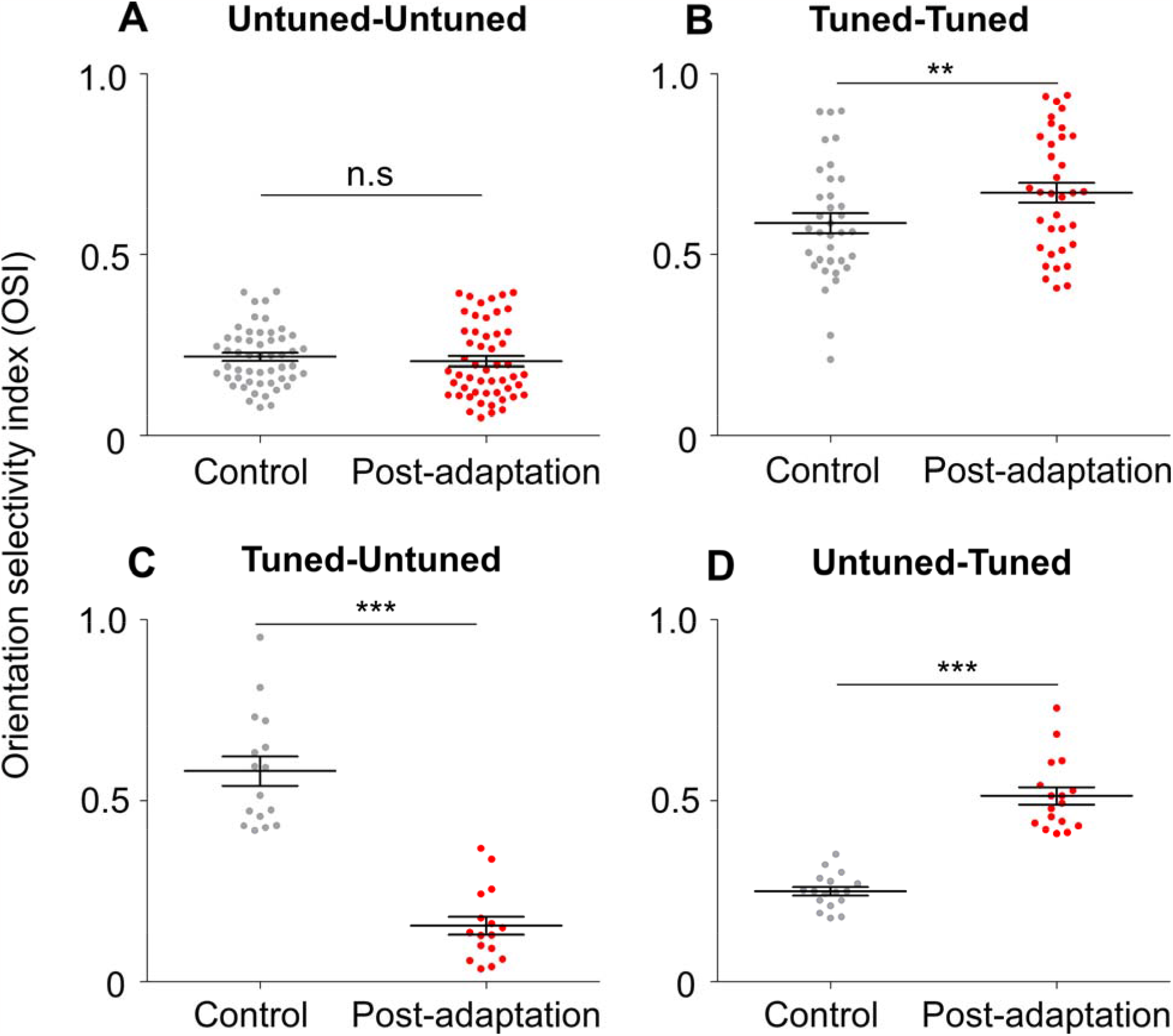
Subpopulation statistics. **(A)** No significant variation between the OSI of the Untuned-Untuned subpopulation in control and post-adaptation conditions (control OSI = 0.21 ± 0.07 vs post-adaptation OSI = 0.20 ± 0.10; Wilcoxon paired test, p = 0.31). **(B)** The OSI of the Tuned-Tuned sub-population increased significantly after adaptation (control OSI = 0.58 ± 0.16 vs post-adaptation OSI = 0.67 ± 0.16; paired t-test, p = 0.009, t = 2.772, df =34). **(C)** Following adaptation, while the OSI of Tuned-Untuned sub-population decreased significantly by 26.52% (control OSI = 0.58 ± 0.15 vs. post-adaptation OSI = 0.15 ± 0.10; paired t-test, p < 0.0001, t = 7.713, df =15), the OSI of the (D) Untuned-Tuned sub-population increased significantly by 48.7% (control OSI = 0.25 ± 0.04 vs post-adaptation OSI = 0.51±0.09; paired t-test, p < 0.0001, t = 9.193, df =16). The bars displayed around the average represent the standard error of mean (SEM).

### 3.2 Correlation of OSI: pre- and post-adaptation

Finally, we hypothesized that adaptation may have distinct effects on the broadly and sharply tuned neurons. To test this, we plotted (**Fig. 4**) OSI post-adaptation as a function of OSI pre-adaptation. We noticed a significant linear deviation from zero (Slope = 0.54 ± 0.10, p < 0.0001) and a modest correlation (Spearman R = 0.40, p < 0.0001) between the cells at the population level (untuned + tuned cells, n = 113), indicating that adaptation further sharpens the selectivity of neurons post-adaptation. This may be because of changed excitation/inhibition balance at the circuit level: oriented inputs at the flanks of the tuning curve may receive enhanced inhibition, resulting in a sharpened tuning curve, thus leading to better learning (Wei et al., 2022).

**Figure 4:**
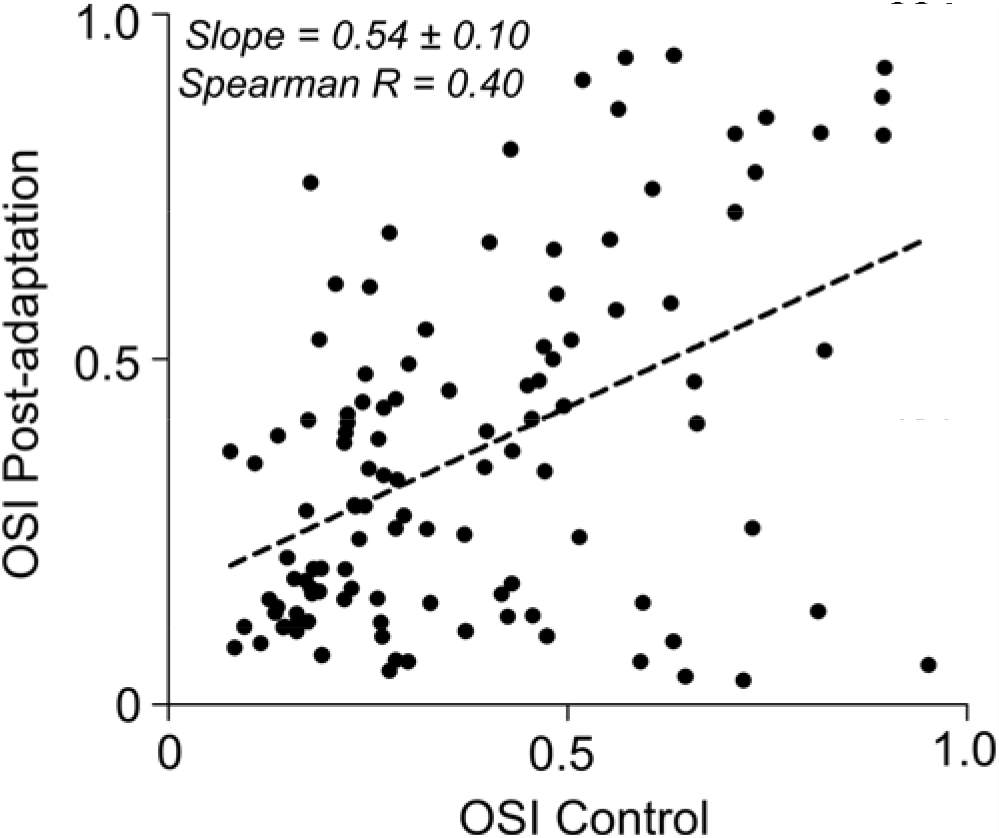
OSI at post-adaptation as a function of OSI at control condition. A modest linear relationship (Slope = 0.54 ± 0.10, Spearman R = 0.40, n = 113) between the conditions suggests that adaptation has more pronounced effect on the highly selective neurons at the control condition.

## 4. DISCUSSION

We investigated the effect of visual adaptation on the mouse visual cortex that has salt-and-pepper organization. We found that post-adaptation a group of cells exhibited significant modification of their OSI following adaptation, whereas another group of neurons maintained their OSI through experimental conditions.

### 4.1. Adaptation sharpens the tuning of visual neurons: homeostasis and E-I balance

Modeling studies have shown that in networks operating in a balanced regime, i.e., where excitation balances inhibition, the tuned component of the input is amplified while the untuned part is suppressed (Hansel and van Vreeswijk, 2012; Sadeh and Rotter, 2015) Consequently, sharp OS, i.e., almost half of units in this study, could emerge in a network even without feature-selective recurrent connectivity. Furthermore, it has also been reported that despite untuned cells having stimulus-independent spiking activity, they contribute to the sensory neural code (Bharmauria et al., 2016a; Pruszynski and Zylberberg, 2019; Safaai et al., 2013). The ratio of tuned and untuned cells along with the average OSI were maintained after twelve minutes of adaptation, suggesting a homeostatic mechanism at the population level (Bachatene et al., 2015a, 2015b; Benucci et al., 2013; Turrigiano and Nelson, 2000).

Feature selectivity, like several cortical functions is affected by the two opposing forces: synaptic excitation and inhibition (Anderson et al., 2000; Carcea and Froemke, 2013; Haider and McCormick, 2009; Sadeh and Clopath, 2021). In our study, the increased selectivity for tuned cells and some untuned cells may be attributed to a shifted excitatory/inhibitory (E-I) balance. For example, in V1 pyramidal cells (Anderson et al., 2000; Haider et al., 2010; Rossi et al., 2019) and other cortical areas (Shu et al., 2003; Wehr and Zador, 2003), increased excitation induced a proportional increase of inhibition, thus stabilizing E/I ratios. This balance can be disrupted by the prolonged viewing of a particular stimulus, the adapter. A computational study reported that a slightly reduced net recurrent cortical excitation could improve the OSI (Teich and Qian, 2003). In addition, it was revealed that adaptation can improve orientation discrimination at the adapted orientation (Regan and Beverley, 1985), and in MT, it affects synaptic inhibition (Qian et al., 1994; Snowden et al., 1991). Thus, we suggest that during the adaptation process, there is a small increase in the strength of recurrent inhibition to cells around the adapted orientation. Indeed, in cortical brain slices, it was shown that repetitive drive leads to a synaptic facilitation between pyramidal and some types of inhibitory neurons (Thomson and Deuchars, 1997). This inhibition could strengthen the degree of selectivity on the flank of the new preferred orientation after adaptation and, thereby, favoring its emergence. A small group of tuned cells showed a decrease of their OSI value. For this group, broadening of tuning curves after adaptation could be explained by a reduction of both net recurrent excitation and inhibition. Indeed, it was demonstrated that, for cells tuned around the adapted orientation, the V1 orientation tuning curves after adaptation became broader (Dragoi et al., 2000) and this can be accounted for by appropriately scaling down both excitation and inhibition to cells around the adapted orientation (Teich and Qian, 2003). This finding strongly suggests that the adaptation induces different changes to the slope of the orientation tuning curves (becoming sharper or broader), that could be explained by an active recalibration of E-I balance in different ways. This might depend on the direction of the shift, toward or away from the adapted orientation or to its amplitude that depends on the cell’s PO in control condition (Bachatene et al., 2013; Ghisovan et al., 2009). Most of the untuned cells remain untuned post-adaptation since their OSI failed to reach statistical significance. Adaptation is predicted to induce changes in the slope of tuning curves; however, we found that it is not the case for most untuned cells. This discrepancy could be explained by different effects of adaptation which would depend on the cell type. It was reported that the tuning curve changes found only occurred in complex cells, whereas simple cells showed no changes at all (Müller et al., 1999).

Neurons work in dedicated circuits to encode a stimulus (Buzsáki, 2010; Molotchnikoff et al., 2019; Singer, 2013). In an ensemble, some connections are flexible and can emerge in the network as the stimulus changes (Bharmauria et al., 2016a; Levy et al., 2020; Singer, 2013). Therefore, we speculate that, following adaptation, stimulus-salient novel connections (Bachatene et al., 2016, 2015a; Bharmauria et al., 2022; Gutnisky and Dragoi, 2008; Kubo et al., 2023) might increase the orientation discrimination of the cell at its post-adaptation PO. Conversely, stable (conserved) connections may not contribute significantly to sharpen the cell’s output (Barth and Poulet, 2012; Molotchnikoff and Rouat, 2012; Rolls and Treves, 2011). In a microcircuit, when a cell fires consistently and connects the same interacting partners pre- and post-adaptation, it generates the same degree of selectivity of the output. However, when a new neuron joins the ensemble, this may affect the cell’s OSI. Briefly, because of the synaptic flexibility of these neuronal groups, a dynamic microcircuit may emerge after adaptation, thus keeping it ever ready to efficiently receive any input output.

### 4.2. Possible molecular mechanisms

Although the molecular mechanisms were not investigated in this study, we speculate that N-methyl-D-aspartate receptor (NMDAR) mechanism might be central to the development / emergence of OSI pre- and post-adaptation. It is well known that orientation discrimination, like most other visual discrimination tasks, is highly sensitive to NMDAR-mediated activity, and plays a critical role in the emergence of OS in V1 (Fox et al., 1989; Miller et al., 1989; Quinlan et al., 1999; Rivadulla et al., 2001; Yu et al., 2008). Notably, when NMDAR antagonists were used, they weakened the amplitude of the orientation map (Ramoa et al., 2001; Yu et al., 2008). In the mouse, repeated presentations of grating stimuli of a single orientation lead to a persistent improvement of the cell’s response, which requires activation of NMDA channels (Frenkel et al., 2006).Thus, when a cell receives new connections due to a repetitive stimulation (adapter), it leads to the increase of NMDAR activation, in turn resulting in changes in calcium flux and other signaling cascades, thereby, sharpening the cell’s response and recentering its tuning around the prevailing stimulus (Kohn and Whitsel, 2002). Moreover, NMDARs are known to be heavily involved in many forms of perceptual learning (Beste et al., 2012; Dinse et al., 2003) and visual adaptation could be considered as a short version of visual learning (Bharmauria et al., 2022; Kohn, 2007; Tring et al., 2023). Thus, adaptation results in short-term facilitation of responses, involving NMDAR, which high-pass-filters orientation selective signals across the synapse (Abbott and Regehr, 2004; Schneggenburger et al., 2002), thus generating highly selective output responses.

## 5. CONCLUSION

All of the above findings underscore that adaptation alters the activity of sharply tuned neurons (putative pyramidal cells), which in turn restructure the entire wiring dynamic of the neuronal assembly by changing the E-I dynamic (Anderson et al., 2000; Rossi et al., 2019). As we did not record the recovery times for neural tuning in this study, thus we cannot speculate about the duration in which such novel tunings are permanently established. However, studies have shown that it takes adaptation log time for neurons to recover (Bharmauria et al., 2022; Tring et al., 2023). Nevertheless, this short-term sharpening of tuning curves may be a central mechanism in learning novel features. Conclusively, it seems that OS is influenced by patterns of neural activity. First, the feedforward thalamo-cortical afferents are related to the establishment of OS (Vidyasagar et al., 1996), then cortical connections play a key role in shaping cortical neuronal tuning properties (Bosking et al., 1997; Mayo and Smith, 2017). This may allow the visual system to support dynamic information processing, thereby, efficiently responding to variations of stimuli.

## Acknowledgements

This study was supported by grants to the Conseil de Recherche en Sciences Naturelles et en Genie du Canada (CRSNG). We also acknowledge Steve Itaya for his comments and writing assistance on the early version of the manuscript. Formatting of funding sources: This work was supported by the Natural Sciences and Engineering Research Council of Canada (NSERC) [RGPIN/04813, 2017].

## Author Contribution

AO and SM conceptualized the study. AO did the experiments and performed the analysis. VB and SM contributed to the data analysis. AO wrote the original draft. VB and SM contributed to writing did the final edits.

## REFERENCES

Abbott LF, Regehr WG (2004) Synaptic computation. Nature 431:796–803.

Afef O, Rudy L, Stéphane M (2022) Ketamine promotes adaption-induced orientation plasticity and vigorous network changes. Brain Res 1797:148111.

Alitto HJ, Dan Y (2010) Function of inhibition in visual cortical processing. Curr Opin Neurobiol 20:340–346.

Anderson JS, Carandini M, Ferster D (2000) Orientation tuning of input conductance, excitation, and inhibition in cat primary visual cortex. J Neurophysiol 84:909–926.

Atallah BV, Bruns W, Carandini M, Scanziani M (2012) Parvalbumin-expressing interneurons linearly transform cortical responses to visual stimuli. Neuron 73:159–170.

Bachatene L, Bharmauria V, Cattan S, Chanauria N, Etindele-Sosso FA, Molotchnikoff S (2016) Functional synchrony and stimulus selectivity of visual cortical units: Comparison between cats and mice. Neuroscience 337.

Bachatene L, Bharmauria V, Cattan S, Chanauria N, Rouat J, Molotchnikoff S (2015a) Summation of connectivity strengths in the visual cortex reveals stability of neuronal microcircuits after plasticity. BMC Neuroscience 16.

Bachatene L, Bharmauria V, Cattan S, Molotchnikoff S (2013) Fluoxetine and serotonin facilitate attractive-adaptation-induced orientation plasticity in adult cat visual cortex. European Journal of Neuroscience 38:2065–2077.

Bachatene L, Bharmauria V, Cattan S, Rouat J, Molotchnikoff S (2015b) Modulation of functional connectivity following visual adaptation: Homeostasis in V1. Brain Research 1594.

Bao M, Fast E, Mesik J, Engel S (2013) Distinct mechanisms control contrast adaptation over different timescales. Journal of Vision 13:14.

Barth AL, Poulet JFA (2012) Experimental evidence for sparse firing in the neocortex. Trends in Neurosciences 35:345–355.

Benucci A, Saleem AB, Carandini M (2013) Adaptation maintains population homeostasis in primary visual cortex. Nat Neurosci 16:724–729.

Beste C, Wascher E, Dinse HR, Saft C (2012) Faster perceptual learning through excitotoxic neurodegeneration. Curr Biol 22:1914–1917.

Bharmauria V, Bachatene L, Cattan S, Brodeur S, Chanauria N, Rouat J, Molotchnikoff S (2016a) Network-selectivity and stimulus-discrimination in the primary visual cortex: Cell-assembly dynamics. European Journal of Neuroscience 43.

Bharmauria V, Bachatene L, Cattan S, Chanauria N, Rouat J, Molotchnikoff S (2015) Stimulus-dependent augmented gamma oscillatory activity between the functionally connected cortical neurons in the primary visual cortex. European Journal of Neuroscience 41:1587–1596.

Bharmauria V, Bachatene L, Ouelhazi A, Cattan S, Chanauria N, Etindele-Sosso FA, Rouat J, Molotchnikoff S (2016b) Interplay of orientation selectivity and the power of low- and high-gamma bands in the cat primary visual cortex. Neuroscience Letters 620.

Bharmauria V, Ouelhazi A, Lussiez R, Molotchnikoff S (2022) Adaptation-induced plasticity in the sensory cortex. J Neurophysiol 128:946–962.

Bonin V, Histed MH, Yurgenson S, Reid RC (2011) Local diversity and fine-scale organization of receptive fields in mouse visual cortex. J Neurosci 31:18506–18521.

Bosking WH, Zhang Y, Schofield B, Fitzpatrick D (1997) Orientation selectivity and the arrangement of horizontal connections in tree shrew striate cortex. J Neurosci 17:2112–2127.

Buzsáki G (2010) Neural syntax: cell assemblies, synapsembles and readers. Neuron 68:362–385.

Campbell FW, Cleland BG, Cooper GF, Enroth-Cugell C (1968) The angular selectivity of visual cortical cells to moving gratings. J Physiol 198:237–250.

Carandini M, Anthony Movshon J, Ferster D (1998) Pattern adaptation and cross-orientation interactions in the primary visual cortex. Neuropharmacology 37:501–511.

Carcea I, Froemke RC (2013) Cortical Plasticity, Excitatory–Inhibitory Balance, and Sensory Perception. Prog Brain Res 207:65–90.

Chanauria N, Bharmauria V, Bachatene L, Cattan S, Rouat J, Molotchnikoff S (2019) Sound Induces Change in Orientation Preference of V1 Neurons: Audio-Visual Cross-Influence. Neuroscience 404:48–61.

Chanauria N, Bharmauria V, Bachatene L, Cattan S, Rouat J, Molotchnikoff S (2016) Comparative effects of adaptation on layers II–III and V–VI neurons in cat V1. European Journal of Neuroscience 44.

Cossell L, Iacaruso MF, Muir DR, Houlton R, Sader EN, Ko H, Hofer SB, Mrsic-Flogel TD (2015) Functional organization of excitatory synaptic strength in primary visual cortex. Nature 518:399–403.

Davey CE, Lloyd EKJ, Kuhlmann L, Burkitt AN, Vidyasagar TR (2022) A neural framework for spontaneous development of orientation selectivity in the primary visual cortex.

Dinse HR, Ragert P, Pleger B, Schwenkreis P, Tegenthoff M (2003) Pharmacological modulation of perceptual learning and associated cortical reorganization. Science 301:91–94.

Dragoi V, Sharma J, Sur M (2000) Adaptation-induced plasticity of orientation tuning in adult visual cortex. Neuron 28:287–98.

Fox K, Sato H, Daw N (1989) The location and function of NMDA receptors in cat and kitten visual cortex. J Neurosci 9:2443–2454.

Frenkel MY, Sawtell NB, Diogo ACM, Yoon B, Neve RL, Bear MF (2006) Instructive Effect of Visual Experience in Mouse Visual Cortex. Neuron 51:339–349.

Ghisovan N, Nemri A, Shumikhina S, Molotchnikoff S (2009) Long adaptation reveals mostly attractive shifts of orientation tuning in cat primary visual cortex. Neuroscience 164:1274–1283.

Giaschi D, Douglas R, Marlin S, Cynader M (1993) The time course of direction-selective adaptation in simple and complex cells in cat striate cortex. Journal of Neurophysiology 70:2024–2034.

Gutnisky DA, Dragoi V (2008) Adaptive coding of visual information in neural populations. Nature 452:220–224.

Haider B, Krause MR, Duque A, Yu Y, Touryan J, Mazer JA, McCormick DA (2010) Synaptic and network mechanisms of sparse and reliable visual cortical activity during nonclassical receptive field stimulation. Neuron 65:107–121.

Haider B, McCormick DA (2009) Rapid neocortical dynamics: cellular and network mechanisms. Neuron 62:171–189.

Hansel D, van Vreeswijk C (2012) The mechanism of orientation selectivity in primary visual cortex without a functional map. J Neurosci 32:4049–4064.

Harris RA, O’Carroll DC, Laughlin SB (2000) Contrast gain reduction in fly motion adaptation. Neuron 28:595–606.

Hubel DH, Wiesel TN (1965) Receptive fields and functional architecture in two nonstriate visual areas (18 and 19) of the cat. Journal of neurophysiology 28:229–289.

Hubel DH, Wiesel TN (1959) Receptive fields of single neurones in the cat’s striate cortex. The Journal of physiology 148:574–91.

Insel N, Barnes CA (2015) Differential Activation of Fast-Spiking and Regular-Firing Neuron Populations During Movement and Reward in the Dorsal Medial Frontal Cortex. Cereb Cortex 25:2631–2647.

Jeyabalaratnam J, Bharmauria V, Bachatene L, Cattan S, Angers A, Molotchnikoff S (2013) Adaptation Shifts Preferred Orientation of Tuning Curve in the Mouse Visual Cortex. PLoS ONE 8.

Kaschube M, Schnabel M, Löwel S, Coppola DM, White LE, Wolf F (2010) Universality in the Evolution of Orientation Columns in the Visual Cortex. Science 330:1113–1116.

Kohn A (2007) Visual adaptation: physiology, mechanisms, and functional benefits. J Neurophysiol 97:3155–3164.

Kohn A, Movshon JA (2004) Adaptation changes the direction tuning of macaque MT neurons. Nature Neuroscience 7:764–772.

Kohn A, Whitsel BL (2002) Sensory cortical dynamics. Behav Brain Res 135:119–126.

Kubo Y, Chalmers E, Luczak A (2023) Biologically-inspired neuronal adaptation improves learning in neural networks. Commun Integr Biol 16:2163131.

Levy M, Sporns O, MacLean JN (2020) Network Analysis of Murine Cortical Dynamics Implicates Untuned Neurons in Visual Stimulus Coding. Cell Reports 31:107483.

Liao DS, Krahe TE, Prusky GT, Medina AE, Ramoa AS (2004) Recovery of cortical binocularity and orientation selectivity after the critical period for ocular dominance plasticity. J Neurophysiol 92:2113–2121.

Mayo JP, Smith MA (2017) Neuronal Adaptation: Tired Neurons or Wired Networks? Trends Neurosci 40:127–128.

Mazurek M, Kager M, Van Hooser SD (2014) Robust quantification of orientation selectivity and direction selectivity. Front Neural Circuits 8:92.

Miller KD, Chapman B, Stryker MP (1989) Visual responses in adult cat visual cortex depend on N-methyl-D-aspartate receptors. Proc Natl Acad Sci U S A 86:5183–5187.

Molotchnikoff S, Bharmauria V, Bachatene L, Chanauria N, Maya-Vetencourt JF (2019) The function of connectomes in encoding sensory stimuli. Progress in Neurobiology 101659.

Molotchnikoff S, Rouat J (2012) Brain at work: time, sparseness and superposition principles. Frontiers in bioscience (Landmark edition) 17:583–606.

Monier C, Chavane F, Baudot P, Graham LJ, Frégnac Y (2003) Orientation and direction selectivity of synaptic inputs in visual cortical neurons: A diversity of combinations produces spike tuning. Neuron 37:663–680.

Müller JR, Metha AB, Krauskopf J, Lennie P (1999) Rapid adaptation in visual cortex to the structure of images. Science 285:1405–1408.

Nguyen G, Freeman AW (2019) A model for the origin and development of visual orientation selectivity. PLoS Comput Biol 15:e1007254.

Niell CM, Stryker MP (2010) Modulation of visual responses by behavioral state in mouse visual cortex. Neuron 65:472–479.

Niell CM, Stryker MP (2008) Highly selective receptive fields in mouse visual cortex. J Neurosci 28:7520–7536.

Ohki K, Chung S, Ch’ng YH, Kara P, Reid RC (2005) Functional imaging with cellular resolution reveals precise micro-architecture in visual cortex. Nature 433:597–603.

Pattadkal JJ, Mato G, van Vreeswijk C, Priebe NJ, Hansel D (2018) Emergent Orientation Selectivity from Random Networks in Mouse Visual Cortex. Cell Rep 24:2042–2050.e6.

Pruszynski JA, Zylberberg J (2019) The language of the brain: real-world neural population codes. Current Opinion in Neurobiology 58:30–36.

Qian N, Andersen RA, Adelson EH (1994) Transparent motion perception as detection of unbalanced motion signals. I. Psychophysics. J Neurosci 14:7357–7366.

Quinlan EM, Philpot BD, Huganir RL, Bear MF (1999) Rapid, experience-dependent expression of synaptic NMDA receptors in visual cortex in vivo. Nature neuroscience 2:352–7.

Ramoa AS, Mower AF, Liao D, Jafri SI (2001) Suppression of cortical NMDA receptor function prevents development of orientation selectivity in the primary visual cortex. J Neurosci 21:4299–4309.

Regan D, Beverley KI (1985) Postadaptation orientation discrimination. J Opt Soc Am A 2:147–155.

Rivadulla C, Sharma J, Sur M (2001) Specific roles of NMDA and AMPA receptors in direction-selective and spatial phase-selective responses in visual cortex. J Neurosci 21:1710–1719.

Roach NW, McGraw PV (2009) Dynamics of spatial distortions reveal multiple time scales of motion adaptation. J Neurophysiol 102:3619–3626.

Rolls ET, Treves A (2011) The neuronal encoding of information in the brain. Progress in Neurobiology 95:448–490.

Rossi LF, Harris KD, Carandini M (2019) Excitatory and inhibitory intracortical circuits for orientation and direction selectivity. bioRxiv 556795.

Sadeh S, Clopath C (2021) Excitatory-inhibitory balance modulates the formation and dynamics of neuronal assemblies in cortical networks. Sci Adv 7:eabg8411.

Sadeh S, Rotter S (2015) Orientation selectivity in inhibition-dominated networks of spiking neurons: effect of single neuron properties and network dynamics. PLoS Comput Biol 11:e1004045.

Safaai H, von Heimendahl M, Sorando JM, Diamond ME, Maravall M (2013) Coordinated population activity underlying texture discrimination in rat barrel cortex. J Neurosci 33:5843–5855.

Schneggenburger R, Sakaba T, Neher E (2002) Vesicle pools and short-term synaptic depression: lessons from a large synapse. Trends Neurosci 25:206–212.

Schneider M, Tzanou A, Uran C, Vinck M (2023) Cell-type-specific propagation of visual flicker. Cell Rep 42:112492.

Shu Y, Hasenstaub A, McCormick DA (2003) Turning on and off recurrent balanced cortical activity. Nature 423:288–293.

Singer W (2013) Cortical dynamics revisited. Trends in Cognitive Sciences 17:616–626.

Snowden RJ, Treue S, Erickson RG, Andersen RA (1991) The response of area MT and V1 neurons to transparent motion. J Neurosci 11:2768–2785.

Swindale NV (1998) Orientation tuning curves: empirical description and estimation of parameters. Biol Cybern 78:45–56.

Tan AYY, Brown BD, Scholl B, Mohanty D, Priebe NJ (2011) Orientation selectivity of synaptic input to neurons in mouse and cat primary visual cortex. The Journal of neuroscience□: the official journal of the Society for Neuroscience 31:12339–50.

Tang MF, Kheradpezhouh E, Lee CCY, Dickinson JE, Mattingley JB, Arabzadeh E (2023) Expectation violations enhance neuronal encoding of sensory information in mouse primary visual cortex. Nat Commun 14:1196.

Teich AF, Qian N (2003) Learning and adaptation in a recurrent model of V1 orientation selectivity. J Neurophysiol 89:2086–2100.

Thomson AM, Deuchars J (1997) Synaptic interactions in neocortical local circuits: dual intracellular recordings in vitro. Cereb Cortex 7:510–522.

Tring E, Dipoppa M, Ringach DL (2023) A power law of cortical adaptation. bioRxiv 2023.05.22.541834.

Turrigiano GG, Nelson SB (2000) Hebb and homeostasis in neuronal plasticity. Current Opinion in Neurobiology 10:358–364.

Vidyasagar TR, Pei X, Volgushev M (1996) Multiple mechanisms underlying the orientation selectivity of visual cortical neurones. Trends Neurosci 19:272–277.

Wehr M, Zador AM (2003) Balanced inhibition underlies tuning and sharpens spike timing in auditory cortex. Nature 426:442–446.

Wei W, Merkt B, Rotter S (2022) A theory of orientation selectivity emerging from randomly sampling the visual field.

Weiler S, Guggiana Nilo D, Bonhoeffer T, Hübener M, Rose T, Scheuss V (2022) Orientation and direction tuning align with dendritic morphology and spatial connectivity in mouse visual cortex. Curr Biol 32:1743–1753.e7.

Wiesel TN, Hubel DH (1963) Single-cell responses in striate cortex of kittens deprived of vision in one eye. Journal of neurophysiology 26:1003–1017.

Wilson DE, Whitney DE, Scholl B, Fitzpatrick D (2016) Orientation selectivity and the functional clustering of synaptic inputs in primary visual cortex. Nat Neurosci 19:1003–1009.

Womelsdorf T, Lima B, Vinck M, Oostenveld R, Singer W, Neuenschwander S, Fries P (2012) Orientation selectivity and noise correlation in awake monkey area V1 are modulated by the gamma cycle. Proc Natl Acad Sci U S A 109:4302–4307.

Yu H, Chen X, Sun C, Shou T (2008) Global evaluation of contributions of GABA A, AMPA and NMDA receptors to orientation maps in cat’s visual cortex. Neuroimage 40:776–787.

